# The skin microbiome as a new potential biomarker in the domestication and welfare of *Octopus vulgaris*

**DOI:** 10.1101/2024.07.11.602832

**Authors:** Daniel Costas-Imbernón, Carolina Costas-Prado, Teresa Sequeiro, Pablo Touriñán, Pablo García-Fernández, Ricardo Tur, David Chavarrías, María Saura, Josep Rotllant

**Affiliations:** Institute of Marine Research-CSIC, Rúa Eduardo Cabello 6, 36208, Vigo, Spain; Pescanova Biomarine Center, Lugar Ardia 172, 36989, O Grove, Spain

**Keywords:** aquaculture, biomarker, common octopus, domestication, dysbiosis, metagenomics, microbiome, welfare

## Abstract

Over the past decade, there has been a growing interest in common octopus aquaculture, prompted by the increasing market demand, the decline in overall fisheries and the search for a more sustainable food resource. Nevertheless, this interest has raised concerns about the potential impact of large-scale production and intensified farming practices in the future. Given the growing interest in octopus aquaculture and society’s increasing concern for animal welfare, this study investigates the mucosal skin microbiota of wild and captive-bred common octopuses. The primary aim is to determine if breeding in captivity affects these communities and consequently animal welfare, using 16S ribosomal RNA metabarcoding. The core microbiota of common octopus mucosal skin is composed of the phyla Bacteroidota, Proteobacteria, Campylobacterota and Verrucomicrobiota, according to our findings. Although variations in abundance were observed, wild and aquaculture octopuses had comparable microbiota composition and diversity. Gammaproteobacteria were found to be enriched in wild mucosal skin samples, and some of the species were described as potentially pathogenic. However, these species were absent or in lower abundance in aquaculture mucosal skin samples. Additionally, analysis of KEGGs predictions showed that wild mucosal skin samples had a higher overall enrichment in functional pathways, primarily associated with xenobiotic remediation pathways, compared to the aquaculture mucosal skin samples. This is the first study to characterize the mucosal skin microbiome of the common octopus and compare the microbiome of wild and aquaculture individuals of this species. The results indicate that current aquaculture practices align with animal welfare through the use of controlled hatchery environments and high-quality water conditions. This study offers valuable insights into the microbiome of the common octopus, which can serve as a biomarker for evaluating animal welfare. It also explores the implications of these findings for the development of sustainable and responsible aquaculture practices.

## 1 Introduction

Aquaculture is a fast-growing industry that plays a significant role as a source of seafood production worldwide, accounting 50% of the total production in 2021 (FAO, 2022). Over the past decade, there has been an increasing interest in cephalopod aquaculture research, driven by the growing market demand, the decline in overall fisheries, and the search for a more sustainable food resource (Berger, 2011; Vidal et al., 2014; Xavier et al., 2015). The common octopus (*Octopus vulgaris*) is considered a promising candidate for aquaculture diversification in Europe, mainly due to its fast growth rate, short life cycle, good adaptability to captive conditions and high nutritional and economic value (Vaz-Pires et al., 2004; Vidal et al., 2014). This offers a solution to meet market demand while also reducing the pressure on wild populations. Objective evaluation of data supports this potential. Despite the significant potential of common octopus aquaculture, its development has been constrained by the complex biological and physiological characteristics of this species, particularly regarding environmental parameters and nutritional requirements (Uriarte et al., 2019, 2011). For years, the high mortality rate in the rearing phase of the paralarvae was the primary bottleneck for aquaculture (Iglesias et al., 2007). The reason for this was nutritional deficiencies, particularly the lack of lipids in the live prey that was frequently utilized as food (Navarro and Villanueva, 2003, 2000). However, this limitation appears to have been resolved with the creation patented rearing technique by scientists from the Spanish Institute of Oceanography and the Pescanova Biomarine Center (Tur et al., 2020). This accomplishment has successfully completed the life cycle of this species, allowing for large-scale industrial cultivation.

The growing interest in octopus aquaculture has also sparked concerns about the future impact of large-scale production and intensified farming practices. In this situation, animal welfare must be a crucial aspect to ensure the development of sustainable and responsible aquaculture practices of this species. Farm animal welfare is generally assessed through measurements of physical health, immune response, behaviour, and physiological indicators, with a focus on identifying stress (Broom, 2010; Fraser et al., 1997). Stress-induced neuroendocrine responses play a critical role in the physiology and overall well-being of animals. While glucocorticoids are recognized as the key hormonal regulators of the physiological stress response in vertebrates (Cockrem, 2013), the underlying mechanisms that govern neuroendocrine responses to stressors in invertebrates, specifically in cephalopods, remain unclear (Fodor and Pirger, 2022).

In recent years, researchers have suggested that the microbial communities present in the host and surrounding environment could serve as a biomarker for health and well-being, owing to their interaction with the host immune system and ability to respond to stressors (Kraimi et al., 2019; Llewellyn et al., 2014; Lorgen-Ritchie et al., 2023). In the case of aquatic animals, the microbial communities on their mucosal surfaces contribute to their individual health by serving a primarily defensive role for the host (Reverter et al., 2018). Previous studies have shown that stress significantly affects the proportion of pathogenic microorganisms on fish skin (Boutin et al., 2013; Chiarello et al., 2015; Mougin and Joyce, 2022; Rich et al., 2023).

Recent advances in high-throughput sequencing technologies and bioinformatics tools have facilitated a more precise investigation of the composition and abundance of microbial communities. Specifically, metagenomics has emerged as the leading technique for analysing genetic material obtained directly from an environmental sample, thereby overcoming the constrains associated whit culture-based methods. These advancements have resulted in a vast scientific literature that has enhanced our comprehension of the interplay between the host, its microbiome and the environment.

Nevertheless, there is still a paucity of research on microbial communities in cephalopods. The literature only includes studies on the gut of common octopus paralarvae by Roura et al., (2017), an analysis of different tissues from *Sepia officinalis* by Lutz et al., (2019), and a comparative investigation of gut microbial composition in six cephalopod species by Kang et al., (2022).

Given the rising interest in octopus aquaculture and the increasing social concern for animal welfare, this study aims to investigate the skin mucus microbial composition on both wild and farmed octopuses. The ultimate goal is to determine whether captive rearing conditions influences these communities and, consequently, animal welfare. Furthermore, we aim to establish the composition of the skin mucus core-microbiota as a tangible biomarker for evaluating and advancing animal welfare in these organisms. To our knowledge, this is the first study to investigate the composition of the microbiota present in the octopus skin mucus of both wild and farmed populations of this species.

## 2 Materials and methods

### 2.1 Sampling

A total of 20 common octopuses of an average weight of approximately 1.2 ± 0.25 kg were sampled from two different sources: wild and aquaculture. The wild octopuses (n = 10, half of each sex) were provided by the San Bartolomé Fishermen’s Guild (Cangas, Galicia), that captured individuals in the Ría of Vigo (42°15′N 8°45′W) in 2022. The farmed octopuses (n = 10) were part of the fifth generation of domestically bred individuals raised and kept at the Pescanova Biomarine Center facilities in O Grove, Spain (42°47′N 8°86′W) in the year 2022. Samples were obtained from 12 m3 square tanks connected to a semi-open water recirculation system, maintaining specific conditions of temperature (15 ± 1 °C), salinity (35 ppm), photoperiod (10 h light:14 h dark), and density (15 kg/m3) (Iglesias, J; Sánchez, F.J.; Otero, J.J.; Moxica, 2000). These tanks were equipped with cognitive enrichment, incorporating diverse structures.

Skin mucus samples were collected from each specimen using sterile cotton swabs. For this purpose, the animals were previously anaesthetised with 1.5% MgCl2 and 1% ethanol. Sampling was performed by gently swabbing along the surface of the octopus. In order to obtain sufficient volumes of skin mucus for downstream analysis, two cotton swabs were taken from the same individual. The swabs were subsequently cut and placed into individual sterile tubes and kept refrigerated on ice until their arrival to the laboratory, where they were frozen at –80 °C until processing.

### 2.2 DNA extraction and sequencing

Microbial DNA extraction was performed following the MasterPure Complete DNA and RNA Purification Kit (Epicentre, Southampton, Hampshire, UK) and Pathogen Lysis Tubes (QIAGEN, Hilden, Germany) protocols, with some modifications proposed by Boix-Amorós et al. (Boix-Amorós et al., 2016) to optimise DNA isolation. Total genomic DNA was extracted from the cotton swabs by adding 500 µL of lysis buffer to each tube and shaking vigorously. The resulting lysates were then transferred to the Pathogen Lysis Tubes and subjected to mechanical disruption using a TissueLyser II (QIAGEN, Hilden, Germany) at 30 Hz for 5 min. After the disruption step, samples were placed on dry ice for 3 min and incubated in a thermoblock at 65 °C for 5 min.

For genomic DNA purification, samples were incubated with 2 µL of proteinase K (50 µg/mL) at 65 °C for 15 min. Next, the tubes were placed on ice to stop the reaction, and 2 µL of RNAse A (5 µg/µL) were added to each sample and incubated at 37 °C for 30 min. After incubation on ice for 3-5 minutes, proteins were then precipitated by adding 250 µL of MPC protein precipitation reagent, followed by vigorous shaking and centrifugation at 10,000 g for 10 minutes at 4 °C. The supernatant was transferred to a clean tube, discarding the pellet. The DNA was precipitated with isopropanol and centrifugated at 4 °C for 10 min at 10,000 g. Finally, the pellet containing the purified DNA was resuspended in 30 µL of TE buffer.

Extracted DNA concentration was measured in a Qubit 4.0 Fluorometer (Thermo Fisher Scientific, DE, United States) using the dsDNA High Sensitivity (HS) Assay Kit (Invitrogen, Carlsbad, California, United States). The purity and quality of the samples were assessed using a NanoDrop 2000 Spectrophotometer (Thermo Fisher Scientific, DE, United States). The microbial community was characterized using the 16S rRNA amplicon sequencing technique, targeting the hypervariable regions V3 and V4. Amplicon library preparation and sequencing were performed by an external service (Foundation for the Promotion of Health and Biomedical Research, FISABIO, Valencia, Spain) using standard protocols. Sequencing was performed using a 2 × 300 bp paired-end protocol on an Illumina MiSeq platform (Illumina, Inc., San Diego, CA, USA). Negative controls were included for both extraction and PCR protocols.

### 2.3 Bioinformatic analyses

Demultiplexed raw reads (FASTQ files) were subjected to quality control with fastp software (Chen et al., 2018) applying a minimum sequence length of 50 nucleotides (*min_length 50*). A quality control step was performed per nucleotide, ensuring that the quality did not fall below 30 (*trim_qual_right 30*) in a 10 nucleotides window (*trim_qual_window 10*), using the mean (*trim_qual_type mean*).

Subsequently, the bioinformatic analysis was conducted following a workflow that includes processing of the reads and taxonomic assignment into ASVs through the QIIME2 v2022.8 pipeline (Bolyen et al., 2019), followed by further statistical analysis using R Statistical Software v4.1.2 (R Core Team, 2013) (Additional file 1: Fig. S1). Filters for abundance or prevalence were not applied. Primer sequences were trimmed from the quality-checked pair-end reads using the *Cutadapt* plugin (Martin, 2011) implemented in QIIME2. Next, data were processed with the DADA2 algorithm (Callahan et al., 2016) to denoise the reads (filtering and chimera removal) and assign them to amplicon sequence variants (ASVs). Taxonomic classification of ASVs was performed using a Naïve Bayes classifier pre-trained on the hypervariable regions V3-V4 with reference sequences extracted from the SILVA (v138) (Quast et al., 2013). Subsequently, sequences assigned to mitochondria and chloroplast, as well as those not assigned at the domain level, were filtered out from the dataset.

### 2.4 Statistical analyses

The ASV table, taxonomy and metadata were imported into R environment using the Phyloseq package v3.17 (McMurdie and Holmes, 2013). The diversity and abundance of the skin microbiota of the wild and aquaculture samples were assessed estimating the alpha and beta diversity.

To conduct alpha diversity analysis, the abundance data were rarefied to the minimum sequencing depth. Alpha diversity metrics, including observed richness, Chao1 index (Chao, 1987), Shannon index (Shannon, 1948) and Inverse Simpson index (Simpson, 1949), were calculated at ASV level using the *estimate_richness* function from Phyloseq. Statistical differences between wild and aquaculture groups, as well as possible differences between sexes were assessed using the non-parametric Wilcoxon rank sum test.

Beta diversity was estimated taking into account the compositional nature of microbiome data (Gloor et al., 2017; Quinn et al., 2018). For this purpose, count data were initially subjected to zero-replacement based on the Count Zero Multiplicative (CZM) method in zCompositions package v1.4.0-1 (Palarea-Albaladejo and Martín-Fernández, 2015) using the default parameters (threshold of 0.5 and frac = 0.65). Next, the CZM adjusted absolute abundances were transformed using the centered log-ratio (CLR) method (Aitchison et al., 2000) with the CoDaSeq package v0.99.6 (Gloor et al., 2016). These values were used to obtain an Euclidean distance matrix (Aitchison distances) that was used to perform a principal component analysis (PCA). Between-group comparisons of beta diversity were assessed using PERMANOVA (Anderson, 2017) with 999 permutations, implemented in the *adonis2* function of the vegan R package v2.6-4 (Dixon, 2003).

To determine differences in the abundance levels of specific taxa, a differential abundance analysis between wild and aquaculture groups was performed with ALDEx2 R package (Fernandes et al., 2014). The input for this analysis was obtained from the CLR-transformed count data generated with the *aldex*.*clr* function using 1,000 Monte Carlo replicates from the Dirichlet distribution. Differences between groups were assessed using Wilcoxon rank sum test through the *aldex*.*ttest* function applying False Discovery Rate (FDR) multitest correction at the 5% level. Values above the significance threshold and with an |effect size factor| (median ratio of between- and within-group differences) ≥ 2.0 were considered for discussion. The results were illustrated in heatmaps, using the ComplexHeatmap R package version 2.14.0 (Gu et al., 2016).

### 2.5 Prediction of functional genes

PICRUSt2 was used to predict potential metabolic functions from 16S rRNA amplicon sequences (Douglas et al., 2020). QIIME2 denoised reads were imported into PICRUSt2 and were compared with sequences from the reference tree. Matches were filtered to exclude those ASV with Nearest Sequenced Taxon Index (the phylogenetic distance between the ASV and the nearest sequenced reference genome) ≥ 2. Functional data were mapped and annotated at different levels of KEGG (Kyoto Encyclopedia of Genes and Genomes) category pathway (Kanehisa et al., 2016). We then used ALDEx2 to identify differentially abundant KEGG between wild and aquaculture sample groups. Pathways were defined as significantly abundant if they met the criteria of an FDR adjusted *p*-value ≤ 0.05 in the Wilcoxon rank sum test and an |effect size| ≥ 1.5.

## 3 Results

After the quality control and pre-processing steps, the final data set corresponding to the 20 skin mucus samples contained 2,283,515 sequences, which were grouped into 6,829 unique ASVs (Additional file 2: Table S1). The mean read counts was 114,175, ranging from 92,046 (sample A4) to 149,578 counts (sample A8). The rarefaction curves revealed that most of bacterial taxa present in the samples were detected, indicating that the sequencing depth was enough to capture most of the bacterial diversity (Additional file 1: Fig. S2).

### 3.1 Community composition

After taxonomic assignment of ASVs, 33 phyla, 74 classes, 203 orders, 350 families and 765 genera were obtained in wild and aquaculture mucosal skin samples.

Of the 33 phyla identified, 24 were common to both groups; however, the prevalence of some phyla varied significantly between samples (Additional file 1: Fig. S3). The most dominant phyla were Bacteroidota, which accounted for 49.6% and 44% of the mean total abundance in wild and aquaculture mucosal skin samples, respectively, followed by Proteobacteria (30.2% and 29.6%), Campylobacterota (6.8% and 10.0%), Verrucomicrobiota (6.9% and 8.9%) and Patescibacteria (2.8% in both groups) (Fig. 1A, B). The relative abundances of these phyla were remarkably similar in both sample types.

**Figure 1.**
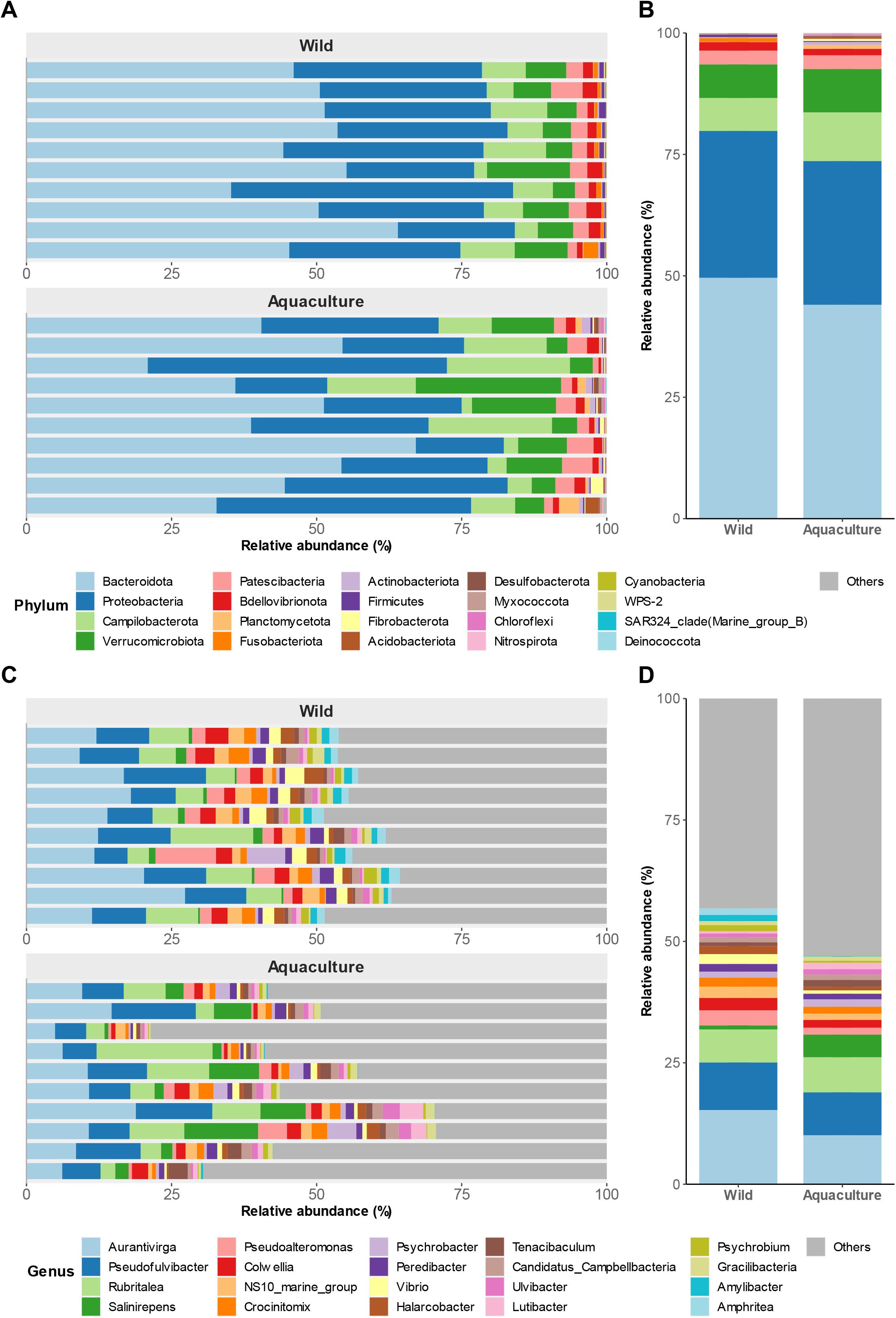
Taxonomic composition of bacterial populations in the skin of wild and aquaculture octopuses. (**A**) Comparation of the relative abundances of the 20 most prevalent phyla (x-axis) found in the mucosal skin microbiome of both wild and aquaculture octopuses. (**B**) Mean relative abundances of these phyla across all samples. **Figures C** and **D** depict the taxonomic composition at the genus level.

At the genus level, no significant differences in the abundance of predominant taxa were observed between the two groups (Fig. 1C, D). Among these genera, *Aurantivirga, Pseudofulvibacter*, and *Rubritalea* proved to be the most abundant, together accounting for approximately 30% of the mean total abundance in wild and aquaculture octopuses. No differences were observed between males and females (data not shown).

### 3.2 Diversity analysis

Results from alpha diversity at the ASVs level revealed significantly higher diversity in wild octopus mucosal skin samples compared to those from aquaculture, as indicated by the Shannon and the Inverse Simpson indexes (*p* ≤ 0.05, Wilcoxon rank sum test), but not by the observed richness and the Chao 1 index (Fig. 2A). These differences disappeared when comparisons were made at the genus level (Additional file 1: Fig. S4A).

**Figure 2.**
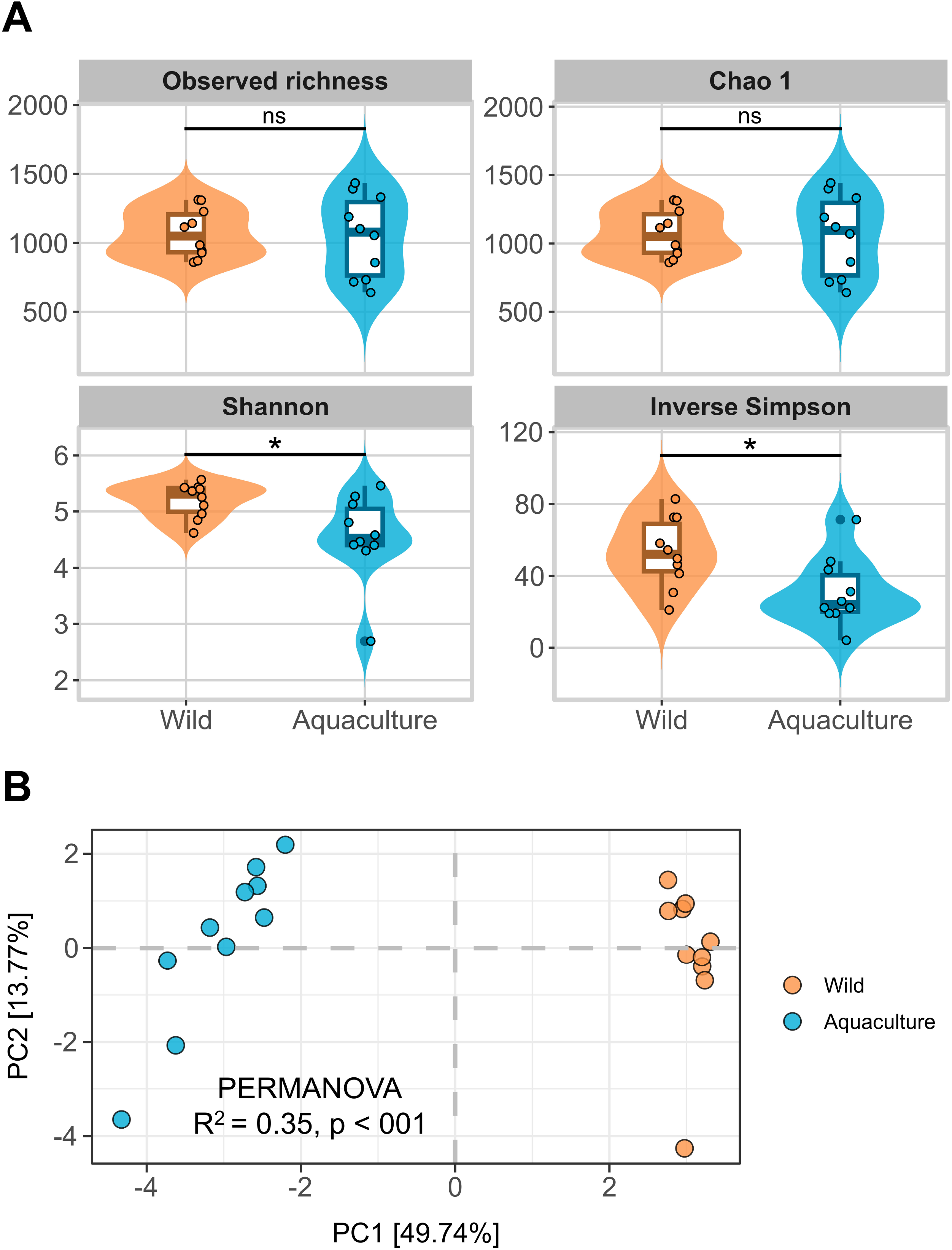
Analysis of diversity. (A) Alpha diversity measures for wild and aquaculture groups are represented by Violin plots that illustrate the alpha diversity indexes analysed (Observed richness, Chao 1, Shannon and Inverse Simpson) at the ASV level in samples from wild and aquaculture groups. The Wilcoxon-Mann-Whitney test results for group comparisons are marked as *ns* (not significant) and asterisk (*p* ≤ 0.05). (B) Beta diversity is analyzed for wild and aquaculture groups. PCA plot displaying Aitchison distances at ASV level. Axes x and y represent the percentage of total variance explained by the first two components, respectively.

Regarding beta diversity, principal component analysis (PCA) on ASV data normalized according to CLR showed a clear distinction between wild and aquaculture octopuses mucosal skin samples, particularly for PC1, which explained 49.7% of the total variance (Fig. 2B). This result was supported by the PERMANOVA test (R^2^ = 0.35, *p* ≤ 0.01), indicating significant differences in the microbial communities between the two groups. When the analysis was performed at the genus level, a greater dispersion was observed in the aquaculture samples (Additional file 1: Fig S4B). Differences between males and females were not significant, as showed by PERMANOVA (R^2^ = 0.12, *p* = 0.239).

### 3.3 Differential abundance

At the species level, from the 500 taxa identified, 80 presented significant differential abundance between wild and aquaculture octopuses mucosal skin samples. Twenty-six out of these 80 species presented additionally an |effect size | ≥ 2.0 (Fig. 3; Additional file 2: Table S2), most of them belonging to the class Gammaproteobacteria. These taxa were predominantly enriched in the skin mucus of wild octopuses (19 in wild vs. 7 in aquaculture). *Tenacibaculum todarodis, Amphritea ceti* and *Pseudoalteromonas marina* were the top enriched (highest size-effect) species in wild samples, while *Flavobacterium frigidarium, Ardenticanaceae sp* and *Chryseobacterium balustinum* were the top enriched species in aquaculture samples. Species previously reported as potentially pathogenic including members from genus *Vibrio* (*V. cortegadensis, V. gallaecicus* and *V. tapetis*), and other taxa such as *Photobacterium swingsii* and *Lactococcus garvieae*, were enriched in the wild group (FDR adjusted *p* ≤ 0.05, Additional file 2: Table S2). At the genus level, the top enriched taxa in wild samples were *Profundimonas, Amphritea* and *Kordia*, while aquaculture samples were enriched in an unidentified genus of Moraxellaceae, *Portibacter* and *Chryseobacterium*. (Additional file 1: Fig. S5; Additional file 2: Table S3).

**Figure 3.**
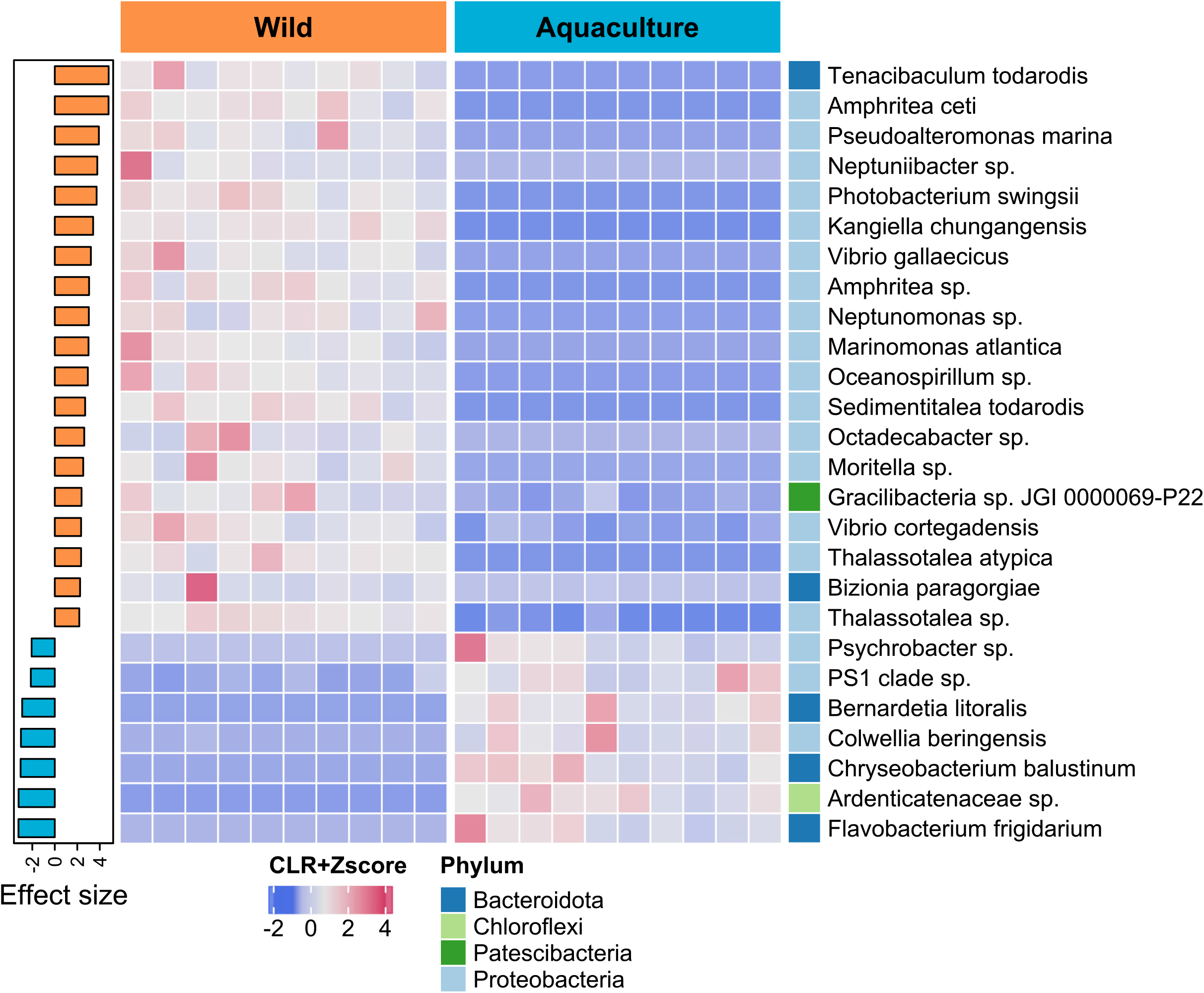
Comparation of abundant species between wild and aquaculture octopuses. The heatmap’s colors represent the scale-transformed relative abundance of 26 bacterial species identified as significantly differentially abundant (FDR adjusted *p* ≤ 0.05 and |effect size| ≥ 2.0). The sample columns are hierarchically clustered, and the bacterial species are ordered by decreasing effect size, as seen in the sidebar plot on the right. The phyla to which each species belongs are also shown on the right sidebar.

### 3.4 Functional profiles of skin microbiomes in wild and farmed octopuses

A total of 7,127 KEGG ortholog (KO) genes were predicted, which were annotated and collapsed into 265 KEGG pathways. From these, 28 pathways were differentially abundant between wild and aquaculture individuals when considering FDR adjusted *p*-values ≤ 0.05 and an |effect size| ≥ 1.5 (Fig. 4). At the highest hierarchy level of classification (level 1), most of the functions were enriched in metabolic pathways, both in wild (12/28 pathways) and aquaculture (4/4 pathways) individuals (Additional file 2: Table S4). In the wild individuals, an enrichment of functions was observed compared to the aquaculture individuals (24 vs 4 functions), several of them associated with the degradation and metabolism of organic compounds such as nitrotoluene, benzoate and geraniol, as well as the biosynthesis of other molecules such as penicillin, cephalosporin and bile acids. On the other hand, the aquaculture individuals presented a higher abundance of pathways related to caffeine metabolism and the biosynthesis of various terpenoids and polyketides.

**Figure 4.**
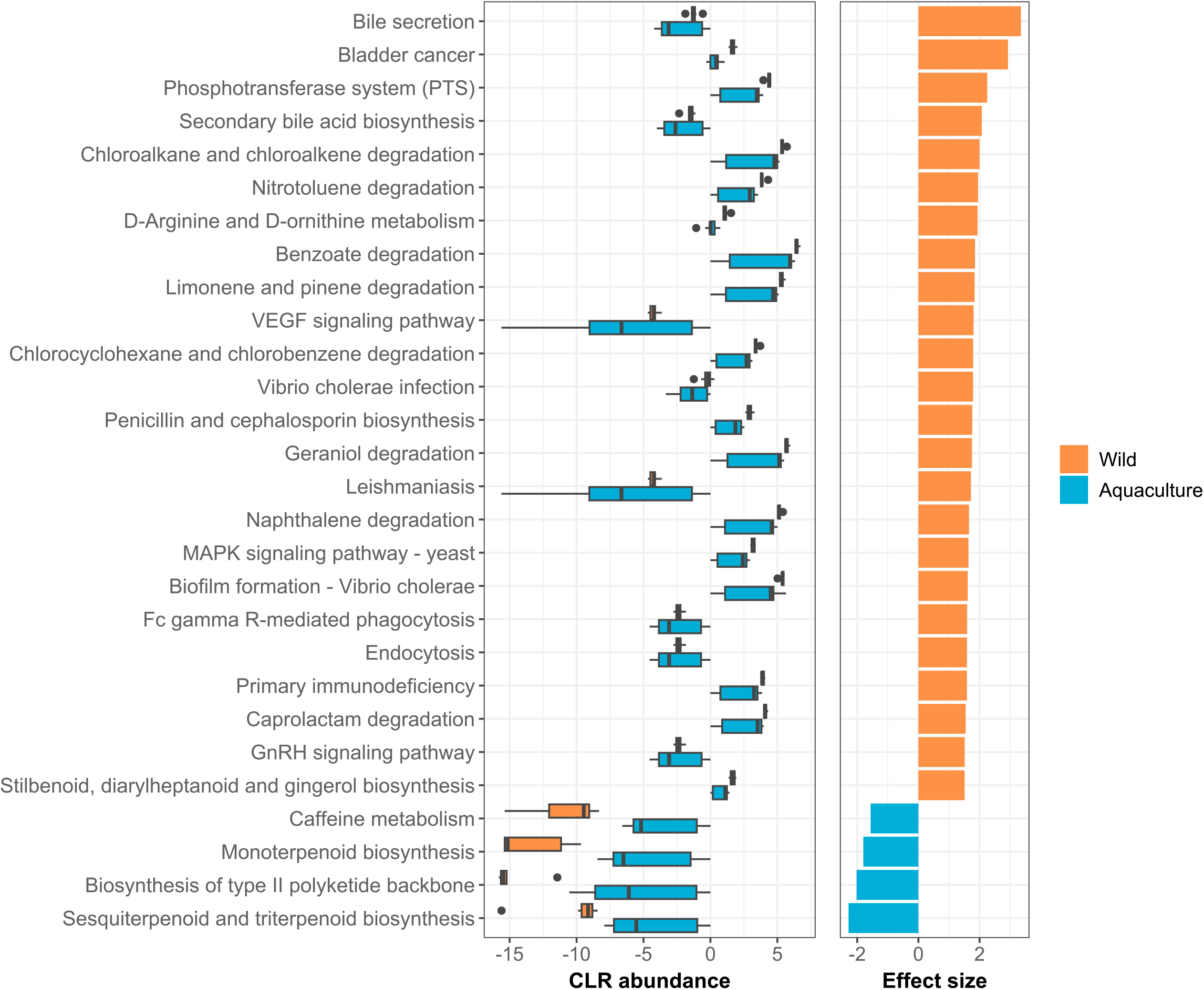
Predicted functional gene profiles. The left panel shows the boxplots of CLR abundances for the 28 PICRUSt2-predicted KEGG pathways identified as differentially abundant between wild and aquaculture groups by ALDEx2 analysis (FDR adjusted *p* ≤ 0.05 and |effect size| ≥ 1.5). Pathways are ordered by descending effect size, shown in the right panel of the graph.

## 4 Discussion

In this study, we performed a comparative analysis of skin mucus microbiome between wild and aquaculture *Octopus vulgaris*. Our results showed that the composition of the microbiota was essentially very similar between the groups of individuals, with differences in beta diversity being mainly attributed to differential abundance. The most relevant difference was related to the presence of bacteria described as pathogenic in wild individuals, which were absent or in lower abundance in aquaculture animals, probably due to the controlled conditions and water quality in the hatchery. To our knowledge, this is the first study where the skin microbiome of the common octopus has been characterized and compared in wild and farmed individuals as a biomarker of animal welfare.

Our results revealed that the microbial communities constituting the common octopus mucosal skin microbiome were dominated by the phylum Bacteroidota, followed by Proteobacteria, Campylobacterota and Verrucomicrobiota. Since these phyla accounted for more than 80% of the abundance in each sample, regardless of their origin, this taxonomic composition can be considered as the core microbiota of the mucosal skin of *Octopus vulgaris*. Although our current understanding of the microbial communities inhabiting cephalopods remains limited, recent studies have begun to shed light on the core microbiota of these animals using high throughput sequencing but focusing primarily on the gut microbiota. Roura et al., (2017) provided an initial overview of the bacterial communities inhabiting the gastrointestinal tract of *O. vulgaris* paralarvae by comparing wild with captive-reared wild individuals using 16S rRNA amplicon sequencing (Roura et al., 2017). In their study, they found that the core gut microbiota was composed of the families Flavobacteriaceae, Comamonadaceae, Moraxellaceae and Sphingomonadaceae, which belong to the phyla Bacteroidota and Proteobacteria, in agreement with our findings. Flavobacteriaceae was also the most abundant family in our study in all samples (Additional file 2: Table S1), followed conversely by Arcobacteraceae and Rubritaleaceae. In contrast to our results, they found important differences between wild and captive-reared wild individuals that they attributed to diet. Lutz et al., (2019) identified a highly simplified microbiota in the skin, gills and gastrointestinal tract of *Sepia officinalis* belonging to *Vibrionaceae* and *Piscirickettsiaceae* families, which were at low frequency or absent in our analysed samples (Lutz et al., 2019). Comparative analysis of six free-living cephalopod species conducted by Kang et al., (2022) revealed that their gut microbiota is composed of distinctive microbes and is strongly associated with the phylogeny of individuals. In their study, they observed that the phyla Tenericutes and Proteobacteria were the most abundant in all samples, while *Mycoplasma* and *Photobacterium* were the core taxa at the genus level, which also differs again from our results. In our study, at the genus level, we observed that *Aurantivirga, Pseudofulvibacter* and *Rubritalea* were the most prevalent, together accounting for about 30% of the total mean abundance in both wild and aquaculture octopuses. Our findings obtained in octopuses differ significantly from the predominance of Proteobacteria over other phyla reported in other marine organisms as described in several fish species (Chiarello et al., 2015; Gomez and Primm, 2021) and other aquatic organisms (Sehnal et al., 2021). Fish skin microbiomes typically include Firmicutes and Actinobacteriota, which in our study showed very low abundances (averaging 0.57 ‐ 0.20% and 0.15 ‐ 0.61%, respectively) in the wild and aquaculture groups. Thus, it is clear that the microbial communities differ markedly depending on the tissue analysed and host genetics.

In general, the alpha diversity was similar in wild and in aquaculture mucosal skin samples. A small discrepancy was found when the analysis was performed at different taxonomic levels. While at the genus level no differences were found between groups for the four indexes estimated, when analysing ASVs, wild samples presented a higher diversity only for those indexes that ignore the taxa evenness and abundance (observed diversity and Chao1 index). However, significance was lost when these factors were taken into account (Shannon and Inverse Simpson indexes), which can be easily explained because filters for relative abundance and prevalence were not applied. This may suggest that, although diversity was similar, the controlled conditions of aquaculture farms, both in terms of diet and prevention for pathogens proliferation may result in a more stable environment and microbiological composition than in wild conditions (Attramadal et al., 2012; Vadstein et al., 2018). Previous studies have evidenced a generalized tendency for alpha diversity to decrease under stress or disease conditions (Legrand et al., 2020; Tarnecki et al., 2019). Along these lines, a loss of evenness and diversity following exposure to stressors has been shown in the microbiome of different marine species (Bagi et al., 2018; Carlson et al., 2017, 2015; He et al., 2017; Narrowe et al., 2015; Nie et al., 2017; Zhang et al., 2018; Zha et al., 2018). Although this is the general trend, changes in diversity are also conditioned by the presence of specific taxa, so a decrease in alpha diversity is not always observed under detrimental conditions (e.g. Vasemägi et al., 2017).

Our results from beta diversity statistics revealed significant differences between wild and aquaculture individuals, which may be due to differences in abundance rather than in the number of taxa detected, according to alpha diversity and differential abundance results. These results could be associated with differences in the environment of wild and aquaculture individuals. Thus, while the wild environment has a heterogeneous nature modulated by different factors, aquaculture specimens were reared in a completely controlled environment, which may influence the mucosal skin microbiome (Lorgen-Ritchie et al., 2021).

Many recent studies have proposed that the microbial patterns found in the microbiota can serve as an indicator of animal health. Thus, by comparing the bacterial composition of healthy and stressed/sick animals, a decrease in beneficial bacteria has been demonstrated, accompanied by an increase in opportunistic bacteria, leading to a decrease in the immune capacity and consequently a higher susceptibility to infections in stressed animals (Legrand et al., 2020; Mougin and Joyce, 2022; Rich et al., 2023). Our findings indicate a widespread decrease in Gammaproteobacteria among aquaculture octopuses. This group of bacteria belongs to the phylum Proteobacteria and encompasses various pathogens, including *Pseudomonas, Salmonella, Vibrio* or *Yersinia* (Maheshwari and Sankar, 2023). Although members of these genera typically inhabit the skin of many marine organisms, some were enriched (by 2.5-fold, Additional file 2: Table S3) in the analyzed samples of wild octopuses.

To investigate whether captive rearing conditions impact welfare in dysbiosis context, we examined the presence and abundance of specific taxa described as potentially pathogenic in aquatic species. Previous studies in cephalopods have reported that pathogenic bacterial infections are caused by gram-negative bacteria belonging to genus *Vibrio* (Fiorito et al., 2015; Gestal et al., 2019; Rich et al., 2023). Our results showed an enrichment of three *Vibrio* species, *V. cortegadensis, V. gallaecicus* and *V. tapetis*, in the wild individuals, although their pathogenicity has not been demonstrated. We also observed an increased abundance of *Photobacterium swingsii* and *Lactococcus garvieae* in the wild individuals. These opportunistic species have been previously isolated from skin lesions in common octopus (Fichi et al., 2015).

The functional prediction analysis revealed that mucosal skin samples from wild octopuses exhibit higher enrichment in metabolic pathways when compare to mucosal skin samples from cultured octopus. The samples from wild octopuses displayed greater enrichment in pathways involving xenobiotic remediation – consistent with their need to adapt to a more diverse and potentially contaminated environment in nature (Jokhakar et al., 2022). The skin microbiome of wild octopus exhibits greater versatility, enabling the degradation of various organic compounds and production of bioactive molecules that have ecological advantages. In contrast, cultured octopuses possess a narrower range of metabolic capacities, which may be influenced by the controlled environment of their cultivation facilities (Lorgen-Ritchie et al., 2021).

While 16S rRNA metabarcoding has limited accuracy in resolving taxa at the species level and making functional prediction (Johnson et al., 2019), it can serve as initial method to obtain details about the taxonomic composition and potential functions of a microbial community (Ortiz-Estrada et al., 2019). The utilization of this method has aided our investigation of the skin mucus microbiome of the octopus as plausible biomarker for animal welfare. Our investigation demonstrated that the managed conditions in farm facilities function as a proactive approach to thwart dysbiosis, underscoring their capacity to uphold the equilibrium of the microbiome and, subsequently, the comprehensive health of farmed octopuses.

## 5 Conclusions

The composition of the common octopus skin microbiota was essentially the same in both wild and aquaculture animals, with a core microbiota dominated by the phyla Bacteroidota, Proteobacteria, Campylobacterota and Verrucomicrobiota. Wild individuals showed a generalized increase in Gammaproteobacteria, including some potential pathogenic bacterial species not found or present in smaller amounts in farmed individuals. In summary, this study offers critical insights into the mucosal skin microbiome of the common octopus as a possible biomarker of animal welfare in this species. Additionally, the study suggests that the controlled conditions in aquaculture facilities may serve as a preventive measure against dysbiosis.

## Data Availability Statement

The datasets generated during the current study are available in the NCBI Sequence Read Archive (SRA) repository under BioProject PRJNA1047176 (https://www.ncbi.nlm.nih.gov/sra/PRJNA1047176).

## Ethics statement

Animal handling was carried out in accordance with the UE welfare directive (Directive 2010/63/UE) and following the authorization file of the animal experimentation project ES360570202001/19/EDUC.FORM.07/JRM.

## Author Contributions

**Daniel Costas-Imbernón:** Conceptualization, Methodology, Validation, Formal analysis, Investigation, Data curation, Writing – original draft, Writing – review and editing, Visualization. **Carolina Costas-Prado:** Validation. **Teresa Sequeiro:** Validation, Formal analysis, Investigation. **Pablo Touriñán:** Resources. **Pablo Gracía-Fernández:** Resources. **Ricardo Tur:** Resources. **David Chavarrías:** Resources. **María Saura:** Conceptualization, Methodology, Formal analysis, Data Curation, Writing – original draft, Writing – review and editing, Visualization, Supervision. **Josep Rotllant:** Conceptualization, Methodology, Resources, Writing – review and editing, Supervision, Project administration, Funding acquisition.

## Acknowledgments

The authors would like to thank Mr. Ruben Chamorro and Ms. Susana Otero for assistance during sample collection.

## Supplementary Material

The Supplementary Material for this article can be found online at: (LINK)

## Supplementary material

### Additional file 1.pdf

**Figure S1**. Bioinformatics workflow used in this study. Raw reads underwent quality control and taxonomic assignment utilizing the SILVA v138 database to generate the ASV table in QIIME2 (left panel). Thereafter, the data were imported into R for statistical analysis using the Phyloseq package (right panel). The analysis focused on taxonomic profiling, with the calculation of alpha and beta diversity, and exploration of differential taxon abundance and potential microbiota functions.

**Figure S2**. Rarefaction curves depicting the number of ASVs as observed richness (y-axis) for wild (orange) and aquaculture (blue) samples are presented, at sequential 1000 read sampling intervals (x-axis).

**Figure S3**. Phyla composition of bacterial populations in the skin of wild and aquaculture octopuses and their prevalence. (A) Venn diagram showing the number of unique and common phyla between both groups. (B) Phyla prevalence. Each dot represents an ASV corresponding to each particular phylum. The average relative abundance is represented on the x-axis while the y-axis displays the proportion of samples in which it is present.

**Figure S4**. Diversity analysis at the genus level. (A) Alpha diversity measures for wild and aquaculture groups. Violin plots represents the alpha diversity indexes analyzed (Observed richness, Chao 1, Shannon and Inverse Simpson) in wild (orange) and aquaculture (blue) samples. (B) Beta diversity for wild and aquaculture groups. PCA plot is based on Aitchison distances at the genus level. Axes x and y represent the percentage of the total variance explained by the first two components, respectively.

**Figure S5**. Differentially abundant genera between wild and aquaculture octopuses. The heatmap colors show the Z-scored CLR-transformed relative abundances of the 37 genera identified as significantly differentially abundant (FDR adjusted *p* ≤ 0.05 and |effect size| ≥ 2.0). The samples are ordered by hierarchical clustering in columns, while the bacterial species are sorted by decreasing effect size in rows, which is illustrated through a side barplot. The right sidebar displays the phyla to which each genus belongs.

### Additional file 2.xlsx

**Table S1**. ASV table and taxonomy. Read counts per sample for all ASVs.

**Table S2**. Results from differential abundance analysis between wild and aquaculture octopuses at species level.

**Table S3**. Results from differential abundance analysis between wild and aquaculture octopuses at genus level.

**Table S4**. Differentially abundant KEGG pathways between wild and aquaculture octopuses.

## References

Aitchison, J., Barceló-Vidal, C., Martín-Fernández, J.A., Pawlowsky-Glahn, V., 2000. Logratio analysis and compositional distance. Math Geol 32, 271–275. 10.1023/A:1007529726302

Anderson, M.J., 2017. Permutational Multivariate Analysis of Variance (PERMANOVA). Wiley StatsRef: Statistics Reference Online 1–15. 10.1002/9781118445112.STAT07841

Attramadal, K.J.K., Salvesen, I., Xue, R., Øie, G., Størseth, T.R., Vadstein, O., Olsen, Y., 2012. Recirculation as a possible microbial control strategy in the production of marine larvae. Aquac Eng 46, 27–39. 10.1016/J.AQUAENG.2011.10.003

Bagi, A., Riiser, E.S., Molland, H.S., Star, B., Haverkamp, T.H.A., Sydnes, M.O., Pampanin, D.M., 2018. Gastrointestinal microbial community changes in Atlantic cod (Gadus morhua) exposed to crude oil. BMC Microbiol 18, 1–14. 10.1186/S12866-018-1171-2/FIGURES/6

Berger, E., 2011. Aquaculture of Octopus species: present status, problems and perspectives. The Plymouth Student Scientist 384–399.

Boix-Amorós, A., Collado, M.C., Mira, A., 2016. Relationship between milk microbiota, bacterial load, macronutrients, and human cells during lactation. Front Microbiol 7, 183883. 10.3389/FMICB.2016.00492/BIBTEX

Bolyen, E., Rideout, J.R., Dillon, M.R., Bokulich, N.A., Abnet, C.C., Al-Ghalith, G.A., Alexander, H., Alm, E.J., Arumugam, M., Asnicar, F., Bai, Y., Bisanz, J.E., Bittinger, K., Brejnrod, A., Brislawn, C.J., Brown, C.T., Callahan, B.J., Caraballo-Rodríguez, A.M., Chase, J., Cope, E.K., Da Silva, R., Diener, C., Dorrestein, P.C., Douglas, G.M., Durall, D.M., Duvallet, C., Edwardson, C.F., Ernst, M., Estaki, M., Fouquier, J., Gauglitz, J.M., Gibbons, S.M., Gibson, D.L., Gonzalez, A., Gorlick, K., Guo, J., Hillmann, B., Holmes, S., Holste, H., Huttenhower, C., Huttley, G.A., Janssen, S., Jarmusch, A.K., Jiang, L., Kaehler, B.D., Kang, K. Bin, Keefe, C.R., Keim, P., Kelley, S.T., Knights, D., Koester, I., Kosciolek, T., Kreps, J., Langille, M.G.I., Lee, J., Ley, R., Liu, Y.X., Loftfield, E., Lozupone, C., Maher, M., Marotz, C., Martin, B.D., McDonald, D., McIver, L.J., Melnik, A. V., Metcalf, J.L., Morgan, S.C., Morton, J.T., Naimey, A.T., Navas-Molina, J.A., Nothias, L.F., Orchanian, S.B., Pearson, T., Peoples, S.L., Petras, D., Preuss, M.L., Pruesse, E., Rasmussen, L.B., Rivers, A., Robeson, M.S., Rosenthal, P., Segata, N., Shaffer, M., Shiffer, A., Sinha, R., Song, S.J., Spear, J.R., Swafford, A.D., Thompson, L.R., Torres, P.J., Trinh, P., Tripathi, A., Turnbaugh, P.J., Ul-Hasan, S., van der Hooft, J.J.J., Vargas, F., Vázquez-Baeza, Y., Vogtmann, E., von Hippel, M., Walters, W., Wan, Y., Wang, M., Warren, J., Weber, K.C., Williamson, C.H.D., Willis, A.D., Xu, Z.Z., Zaneveld, J.R., Zhang, Y., Zhu, Q., Knight, R., Caporaso, J.G., 2019. Reproducible, interactive, scalable and extensible microbiome data science using QIIME 2. Nat. Biotech. 2019 37:8 37, 852–857. 10.1038/s41587-019-0209-9

Boutin, S., Bernatchez, L., Audet, C., Derôme, N., 2013. Network Analysis Highlights Complex Interactions between Pathogen, Host and Commensal Microbiota. PLoS One 8, e84772. 10.1371/JOURNAL.PONE.0084772

Broom, D.M., 2010. Animal welfare: An aspect of care, sustainability, and food quality required by the public. J Vet Med Educ 37, 83–88. 10.3138/jvme.37.1.83

Callahan, B.J., McMurdie, P.J., Rosen, M.J., Han, A.W., Johnson, A.J.A., Holmes, S.P., 2016. DADA2: High-resolution sample inference from Illumina amplicon data. Nat Methods 2016 13:7 13, 581–583. 10.1038/nmeth.3869

Carlson, J.M., Hyde, E.R., Petrosino, J.F., Manage, A.B.W., Primm, T.P., 2015. The host effects of Gambusia affinis with an antibiotic-disrupted microbiome. Comp Biochem Physiol C Toxicol Pharmacol 178, 163–168. 10.1016/J.CBPC.2015.10.004

Carlson, J.M., Leonard, A.B., Hyde, E.R., Petrosino, J.F., Primm, T.P., 2017. Microbiome disruption and recovery in the fish <em>Gambusia affinis</em> following exposure to broad-spectrum antibiotic. Infect Drug Resist 10, 143–154. 10.2147/IDR.S129055

Chao, A., 1987. Estimating the Population Size for Capture-Recapture Data with Unequal Catchability. Biometrics 43, 783. 10.2307/2531532

Chen, S., Zhou, Y., Chen, Y., Gu, J., 2018. fastp: an ultra-fast all-in-one FASTQ preprocessor. Bioinformatics 34, i884–i890. 10.1093/BIOINFORMATICS/BTY560

Chiarello, M., Villéger, S., Bouvier, C., Bettarel, Y., Bouvier, T., 2015. High diversity of skin-associated bacterial communities of marine fishes is promoted by their high variability among body parts, individuals and species. FEMS Microbiol Ecol 91. 10.1093/FEMSEC/FIV061

Cockrem, J.F., 2013. Individual variation in glucocorticoid stress responses in animals. Gen Comp Endocrinol 181, 45–58. 10.1016/J.YGCEN.2012.11.025

Dixon, P., 2003. VEGAN, a package of R functions for community ecology. J VEG SCI 14, 927–930. 10.1111/J.1654-1103.2003.TB02228.X

Douglas, G.M., Maffei, V.J., Zaneveld, J.R., Yurgel, S.N., Brown, J.R., Taylor, C.M., Huttenhower, C., Langille, M.G.I., 2020. PICRUSt2 for prediction of metagenome functions. Nat Biotech 2020 38:6 38, 685–688. 10.1038/s41587-020-0548-6

FAO, 2022. The State of World Fisheries and Aquaculture 2022, The State of World Fisheries and Aquaculture 2022. FAO. 10.4060/cc0461en

Fernandes, A.D., Reid, J.N.S., Macklaim, J.M., McMurrough, T.A., Edgell, D.R., Gloor, G.B., 2014. Unifying the analysis of high-throughput sequencing datasets: Characterizing RNA-seq, 16S rRNA gene sequencing and selective growth experiments by compositional data analysis. Microbiome 2, 1–13. 10.1186/2049-2618-2-15/COMMENTS

Fichi, G., Cardeti, G., Perrucci, S., Vanni, A., Cersini, A., Lenzi, C., De Wolf, T., Fronte, B., Guarducci, M., Susini, F., 2015. Skin lesion-associated pathogens from Octopus vulgaris: first detection of Photobacterium swingsii, Lactococcus garvieae and betanodavirus. Dis Aquat Organ 115, 147–156. 10.3354/DAO02877

Fiorito, G., Affuso, A., Basil, J., Cole, A., de Girolamo, P., D’angelo, L., Dickel, L., Gestal, C., Grasso, F., Kuba, M., Mark, F., Melillo, D., Osorio, D., Perkins, K., Ponte, G., Shashar, N., Smith, D., Smith, J., Andrews, P. lr, 2015. Guidelines for the Care and Welfare of Cephalopods in Research –A consensus based on an initiative by CephRes, FELASA and the Boyd Group. Lab Anim 49, 1–90. 10.1177/0023677215580006/ASSET/IMAGES/LARGE/10.1177_0023677215580006-FIG1.JPEG

Fodor, I., Pirger, Z., 2022. From Dark to Light – An Overview of Over 70 Years of Endocrine Disruption Research on Marine Mollusks. Front Endocrinol (Lausanne) 13, 903575. 10.3389/FENDO.2022.903575/BIBTEX

Fraser, D., Weary, D.M., Pajor, E.A., Milligan, B.N., 1997. A Scientific Conception of Animal Welfare that Reflects Ethical Concerns. Animal Welfare 6, 187–205. 10.1017/S0962728600019795

Gestal, C., Pascual, S., Guerra, Á., Fiorito, G., Vieites Editors, J.M., 2019. Handbook of Pathogens and Diseases in Cephalopods. Handbook of Pathogens and Diseases in Cephalopods 230. 10.1007/978-3-030-11330-8

Gloor, G.B., Macklaim, J.M., Pawlowsky-Glahn, V., Egozcue, J.J., 2017. Microbiome datasets are compositional: And this is not optional. Front Microbiol 8, 2224. 10.3389/FMICB.2017.02224/BIBTEX

Gloor, G.B., Wu, J.R., Pawlowsky-Glahn, V., Egozcue, J.J., 2016. It’s all relative: analyzing microbiome data as compositions. Ann Epidemiol 26, 322–329. 10.1016/J.ANNEPIDEM.2016.03.003

Gomez, J.A., Primm, T.P., 2021. A Slimy Business: the Future of Fish Skin Microbiome Studies. Microb Ecol 82, 275–287. 10.1007/S00248-020-01648-W/TABLES/2

Gu, Z., Eils, R., Schlesner, M., 2016. Complex heatmaps reveal patterns and correlations in multidimensional genomic data. Bioinformatics 32, 2847–2849. 10.1093/BIOINFORMATICS/BTW313

He, S., Wang, Q., Li, S., Ran, C., Guo, X., Zhang, Z., Zhou, Z., 2017. Antibiotic growth promoter olaquindox increases pathogen susceptibility in fish by inducing gut microbiota dysbiosis. Sci China Life Sci 60, 1260–1270. 10.1007/S11427-016-9072-6/METRICS

Iglesias, J., Sánchez, F.J., Bersano, J.G.F., Carrasco, J.F., Dhont, J., Fuentes, L., Linares, F., Muñoz, J.L., Okumura, S., Roo, J., van der Meeren, T., Vidal, E.A.G., Villanueva, R., 2007. Rearing of Octopus vulgaris paralarvae: Present status, bottlenecks and trends. Aquaculture 266, 1–15. 10.1016/J.AQUACULTURE.2007.02.019

Iglesias, J; Sánchez, F.J.; Otero, J.J.; Moxica, C., 2000. Culture of octopus (Octopus vulgaris, Cuvier): Present knowledge, problems and perspectives. Cah. Options Méditerr 47, 313–321.

Johnson, J.S., Spakowicz, D.J., Hong, B.Y., Petersen, L.M., Demkowicz, P., Chen, L., Leopold, S.R., Hanson, B.M., Agresta, H.O., Gerstein, M., Sodergren, E., Weinstock, G.M., 2019. Evaluation of 16S rRNA gene sequencing for species and strain-level microbiome analysis. Nat Commun 2019 10:1 10, 1–11. 10.1038/s41467-019-13036-1

Jokhakar, P., Godhaniya, M., Vaghamshi, N., Patel, R., Ghelani, A., Dudhagara, P., 2022. Comparative taxonomic and functional microbiome profiling of anthrospheric river tributary for xenobiotics degradation study. Ecol Genet Genom 25, 100144. 10.1016/J.EGG.2022.100144

Kanehisa, M., Sato, Y., Kawashima, M., Furumichi, M., Tanabe, M., 2016. KEGG as a reference resource for gene and protein annotation. Nucleic Acids Res 44, D457–D462. 10.1093/NAR/GKV1070

Kang, W., Kim, P.S., Tak, E.J., Sung, H., Shin, N.-R., Hyun, D.-W., Whon, T.W., Kim, H.S., Lee, J.-Y., Yun, J.-H., Jung, M.-J., Bae, J.-W., 2022. Host phylogeny, habitat, and diet are main drivers of the cephalopod and mollusk gut microbiome. Animal Microbiome 2022 4:1 4, 1–15. 10.1186/S42523-022-00184-X

Kraimi, N., Dawkins, M., Gebhardt-Henrich, S.G., Velge, P., Rychlik, I., Volf, J., Creach, P., Smith, A., Colles, F., Leterrier, C., 2019. Influence of the microbiota-gut-brain axis on behavior and welfare in farm animals: A review. Physiol Behav 210, 112658. 10.1016/J.PHYSBEH.2019.112658

Legrand, T.P.R.A., Wynne, J.W., Weyrich, L.S., Oxley, A.P.A., 2020. A microbial sea of possibilities: current knowledge and prospects for an improved understanding of the fish microbiome. Rev Aquac 12, 1101–1134. 10.1111/RAQ.12375

Llewellyn, M.S., Boutin, S., Hoseinifar, S.H., Derome, N., 2014. Teleost microbiomes: The state of the art in their characterization, manipulation and importance in aquaculture and fisheries. Front Microbiol 5, 1–1. 10.3389/FMICB.2014.00207/BIBTEX

Lorgen-Ritchie, M., Clarkson, M., Chalmers, L., Taylor, J.F., Migaud, H., Martin, S.A.M., 2021. A Temporally Dynamic Gut Microbiome in Atlantic Salmon During Freshwater Recirculating Aquaculture System (RAS) Production and Post-seawater Transfer. Front Mar Sci 8, 869. 10.3389/FMARS.2021.711797/BIBTEX

Lorgen-Ritchie, M., Uren Webster, T., McMurtrie, J., Bass, D., Tyler, C.R., Rowley, A., Martin, S.A.M., 2023. Microbiomes in the context of developing sustainable intensified aquaculture. Front Microbiol 14, 1200997. 10.3389/FMICB.2023.1200997/BIBTEX

Lutz, H.L., Ramírez-Puebla, S.T., Abbo, L., Durand, A., Schlundt, C., Gottel, N.R., Sjaarda, A.K., Hanlon, R.T., Gilbert, J.A., Mark Welch, J.L., 2019. A Simple Microbiome in the European Common Cuttlefish, Sepia officinalis. mSystems 4. 10.1128/msystems.00177-19

Maheshwari, P., Sankar, P.M., 2023. Culture-independent and culture-dependent approaches in symbiont analysis: in proteobacteria. Microbial Symbionts: Functions and Molecular Interactions on Host 743–763. Academic Press. 10.1016/B978-0-323-99334-0.00018-9

Martin, M., 2011. Cutadapt removes adapter sequences from high-throughput sequencing reads. EMBnet J 17, 10–12. 10.14806/ej.17.1.200

McMurdie, P.J., Holmes, S., 2013. phyloseq: An R Package for Reproducible Interactive Analysis and Graphics of Microbiome Census Data. PLoS One 8, e61217. 10.1371/JOURNAL.PONE.0061217

Mougin, J., Joyce, A., 2022. Fish disease prevention via microbial dysbiosis-associated biomarkers in aquaculture. Rev Aquac. 10.1111/RAQ.12745

Narrowe, A.B., Albuthi-Lantz, M., Smith, E.P., Bower, K.J., Roane, T.M., Vajda, A.M., Miller, C.S., 2015. Perturbation and restoration of the fathead minnow gut microbiome after low-level triclosan exposure. Microbiome 3, 1–18. 10.1186/s40168-015-0069-6

Navarro, J.C., Villanueva, R., 2003. The fatty acid composition of Octopus vulgaris paralarvae reared with live and inert food: deviation from their natural fatty acid profile. Aquaculture 219, 613–631. 10.1016/S0044-8486(02)00311-3

Navarro, J.C., Villanueva, R., 2000. Lipid and fatty acid composition of early stages of cephalopods: an approach to their lipid requirements. Aquaculture 183, 161–177. 10.1016/S0044-8486(99)00290-2

Nie, L., Zhou, Q.J., Qiao, Y., Chen, J., 2017. Interplay between the gut microbiota and immune responses of ayu (Plecoglossus altivelis) during Vibrio anguillarum infection. Fish Shellfish Immunol 68, 479–487. 10.1016/J.FSI.2017.07.054

Ortiz-Estrada, Á.M., Gollas-Galván, T., Martínez-Córdova, L.R., Martínez-Porchas, M., 2019. Predictive functional profiles using metagenomic 16S rRNA data: a novel approach to understanding the microbial ecology of aquaculture systems. Rev Aquac 11, 234–245. 10.1111/RAQ.12237

Palarea-Albaladejo, J., Martín-Fernández, J.A., 2015. zCompositions — R package for multivariate imputation of left-censored data under a compositional approach. Chemometrics and Intelligent Laboratory Systems 143, 85–96. 10.1016/J.CHEMOLAB.2015.02.019

Quast, C., Pruesse, E., Yilmaz, P., Gerken, J., Schweer, T., Yarza, P., Peplies, J., Glöckner, F.O., 2013. The SILVA ribosomal RNA gene database project: improved data processing and web-based tools. Nucleic Acids Res 41, D590–D596. 10.1093/NAR/GKS1219

Quinn, T.P., Erb, I., Richardson, M.F., Crowley, T.M., 2018. Understanding sequencing data as compositions: an outlook and review. Bioinformatics 34, 2870–2878. 10.1093/BIOINFORMATICS/BTY175

R Core Team, 2013. R: A language and environment for statistical computing.

Reverter, M., Tapissier-Bontemps, N., Lecchini, D., Banaigs, B., Sasal, P., 2018. Biological and Ecological Roles of External Fish Mucus: A Review. Fishes 2018, Vol. 3, Page 41 3, 41. 10.3390/FISHES3040041

Rich, A.F., Denk, D., Sangster, C.R., Stidworthy, M.F., 2023. A retrospective study of pathologic findings in cephalopods (extant subclasses: Coleoidea and Nautiloidea) under laboratory and aquarium management. Vet Pathol 578–598. 10.1177/03009858231186306

Roura, Á., Doyle, S.R., Nande, M., Strugnell, J.M., 2017. You are what you eat: A genomic analysis of the gut microbiome of captive and wild Octopus vulgaris Paralarvae and their zooplankton prey. Front Physiol 8, 362. 10.3389/FPHYS.2017.00362/BIBTEX

Sehnal, L., Brammer-Robbins, E., Wormington, A.M., Blaha, L., Bisesi, J., Larkin, I., Martyniuk, C.J., Simonin, M., Adamovsky, O., 2021. Microbiome Composition and Function in Aquatic Vertebrates: Small Organisms Making Big Impacts on Aquatic Animal Health. Front Microbiol 12, 567408. 10.3389/FMICB.2021.567408/BIBTEX

Shannon, C.E., 1948. A Mathematical Theory of Communication. Bell System Technical Journal 27, 379–423. 10.1002/J.1538-7305.1948.TB01338.X

Simpson, E.H., 1949. Measurement of Diversity. Nature 1949 163:4148 163, 688–688. 10.1038/163688a0

Tarnecki, A.M., Brennan, N.P., Schloesser, R.W., Rhody, N.R., 2019. Shifts in the Skin-Associated Microbiota of Hatchery-Reared Common Snook Centropomus undecimalis During Acclimation to the Wild. Microb Ecol 77, 770–781. 10.1007/s00248-018-1252-7

Tur, R., Rodrigues dos santos, P., Almansa, E., Lago, M., Garcia, P., Perez, E., 2020. Procedimiento para el cultivo de paralarvas del pulpo común Octopus vulgaris-Instituto Español de Oceanografía.

Uriarte, I., Astorga, M., Navarro, J.C., Viana, M.T., Rosas, C., Molinet, C., Hernández, J., Navarro, J., Moreno-Villoslada, I., Amthauer, R., Kausel, G., Figueroa, J., Paredes, E., Paschke, K., Romero, A., Hontoria, F., Varó, I., Vargas-Chacoff, L., Toro, J., Yáñez, A., Cárdenas, L., Enriquez, R., Olivares, A., Rey, M., Izquierdo, M., Sorgeloos, P., Soto, D., Farías, A., 2019. Early life stage bottlenecks of carnivorous molluscs under captivity: a challenge for their farming and contribution to seafood production. Rev Aquac 11, 431–457. 10.1111/RAQ.12240

Uriarte, I., Iglesias, J., Domingues, P., Rosas, C., Viana, M.T., Navarro, J.C., Seixas, P., Vidal, E., Ausburger, A., Pereda, S., Godoy, F., Paschke, K., Farías, A., Olivares, A., Zuñiga, O., 2011. Current Status and Bottle Neck of Octopod Aquaculture: The Case of American Species. J World Aquac Soc 42, 735–752. 10.1111/J.1749-7345.2011.00524.X

Vadstein, O., Attramadal, K.J.K., Bakke, I., Olsen, Y., 2018. K-selection as microbial community management strategy: A method for improved viability of larvae in aquaculture. Front Microbiol 9, 367702. 10.3389/FMICB.2018.02730/BIBTEX

Vasemägi, A., Visse, M., Kisand, V., 2017. Effect of Environmental Factors and an Emerging Parasitic Disease on Gut Microbiome of Wild Salmonid Fish. mSphere 2. 10.1128/msphere.00418-17

Vaz-Pires, P., Seixas, P., Barbosa, A., 2004. Aquaculture potential of the common octopus (Octopus vulgaris Cuvier, 1797): a review. Aquaculture 238, 221–238. 10.1016/J.AQUACULTURE.2004.05.018

Vidal, E.A.G., Villanueva, R., Andrade, J.P., Gleadall, I.G., Iglesias, J., Koueta, N., Rosas, C., Segawa, S., Grasse, B., Franco-Santos, R.M., Albertin, C.B., Caamal-Monsreal, C., Chimal, M.E., Edsinger-Gonzales, E., Gallardo, P., Le Pabic, C., Pascual, C., Roumbedakis, K., Wood, J., 2014. Cephalopod Culture: Current Status of Main Biological Models and Research Priorities. Adv Mar Biol 67, 1–98. 10.1016/B978-0-12-800287-2.00001-9

Xavier, J.C., Allcock, A.L., Cherel, Y., Lipinski, M.R., Pierce, G.J., Rodhouse, P.G.K., Rosa, R., Shea, E.K., Strugnell, J.M., Vidal, E.A.G., Villanueva, R., Ziegler, A., 2015. Future challenges in cephalopod research*. J Mar Biol Assoc U. K. 95, 999–1015. 10.1017/S0025315414000782

Zhang, Xiaoting, Ding, L., Yu, Y., Kong, W., Yin, Y., Huang, Z., Zhang, Xuezhen, Xu, Z., 2018. The Change of Teleost Skin Commensal Microbiota Is Associated With Skin Mucosal Transcriptomic Responses During Parasitic Infection by Ichthyophthirius multifillis. Front Immunol 9, 416680. 10.3389/FIMMU.2018.02972/BIBTEX

Zha, Y., Eiler, A., Johansson, F., Svanbäck, R., 2018. Effects of predation stress and food ration on perch gut microbiota. Microbiome 6, 1–12. 10.1186%2Fs40168-018-0400-0

