## Additional file 1 for "The skin microbiome as a new potential biomarker in the domestication and welfare of *Octopus vulgaris*"

### Supplementary information

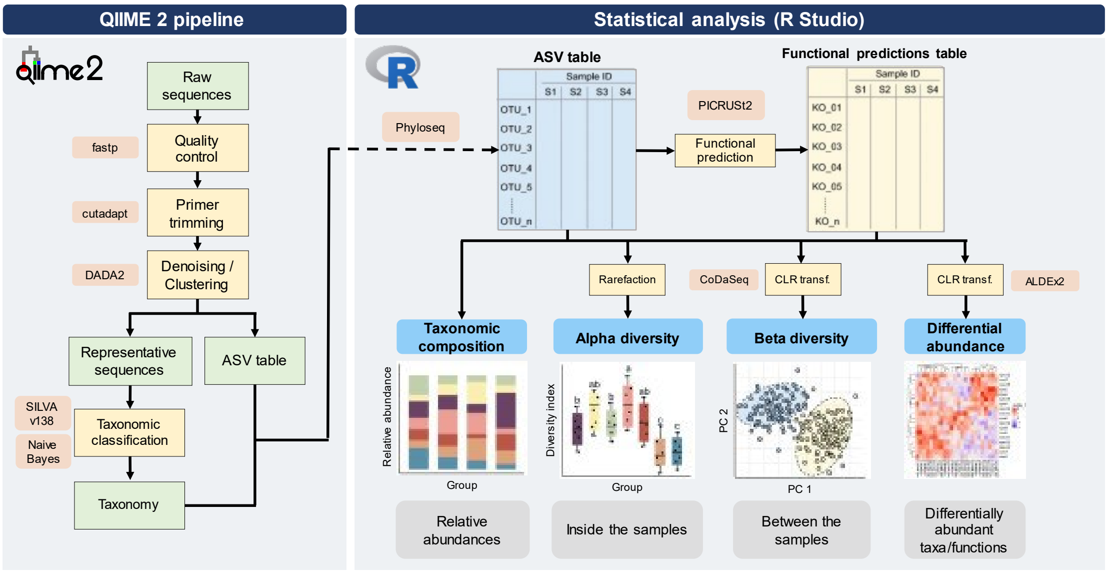


**Figure S1.** Bioinformatics workflow used in this study. Raw reads underwent quality control and taxonomic assignment utilizing the SILVA v138 database to generate the ASV table in QIIME2 (left panel). Thereafter, the data were imported into R for statistical analysis using the Phyloseq package (right panel). The analysis focused on taxonomic profiling, with the calculation of alpha and beta diversity, and exploration of differential taxon abundance and potential microbiota functions.

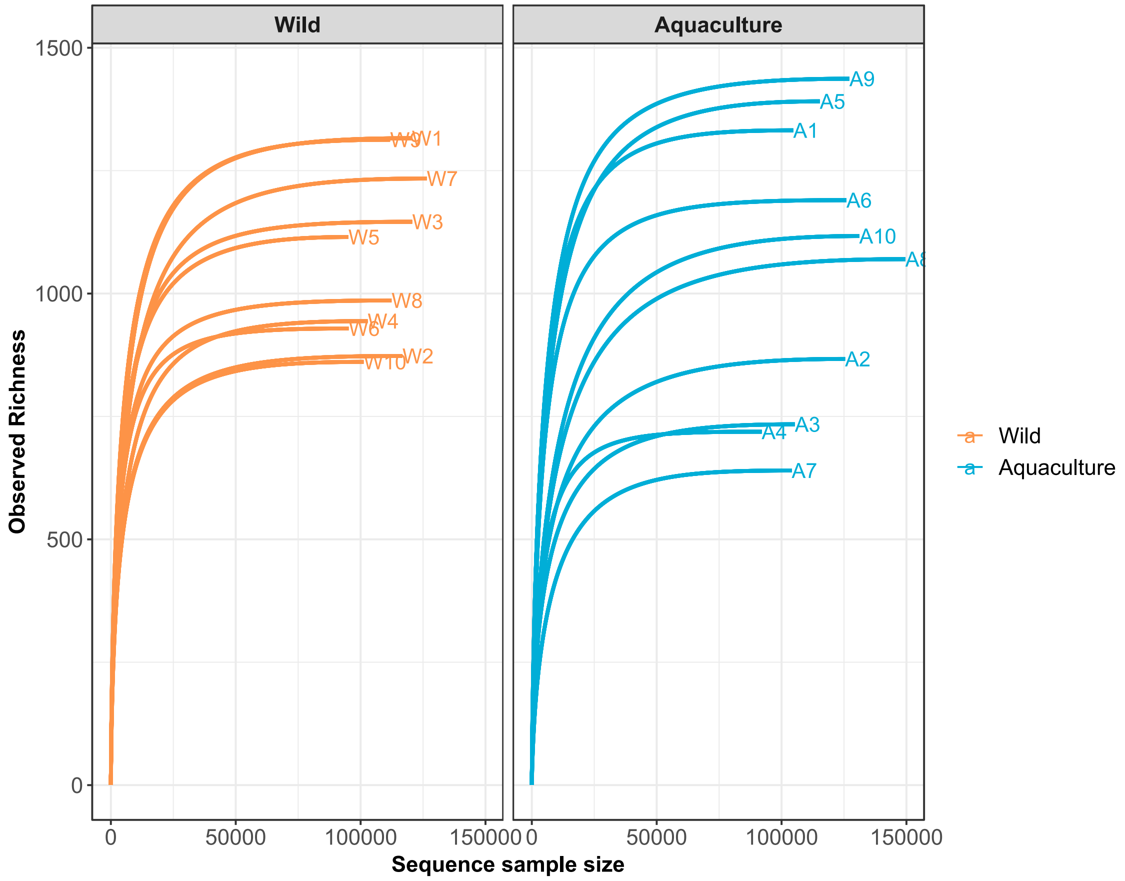


**Figure S2.** Rarefaction curves depicting the number of ASVs as observed richness (y-axis) for wild (orange) and aquaculture (blue) samples are presented, at sequential 1000 read sampling intervals (x-axis).

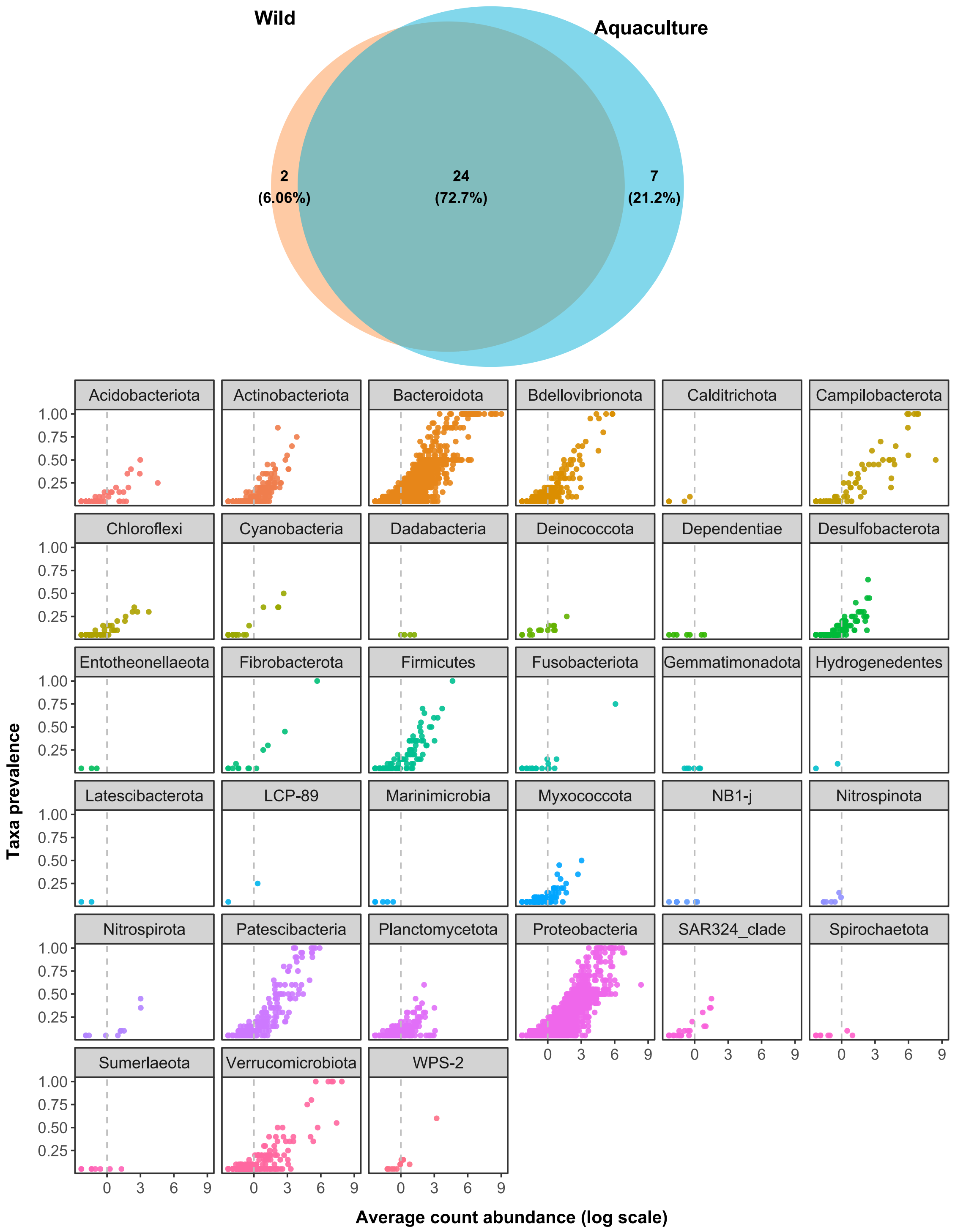


**Figure S3.** Phyla composition of bacterial populations in the skin of wild and aquaculture octopuses and their prevalence. (A) Venn diagram showing the number of unique and common phyla between both groups. (B) Phyla prevalence. Each dot represents an ASV corresponding to each particular phylum. The average relative abundance is represented on the x-axis while the y-axis displays the proportion of samples in which it is present.

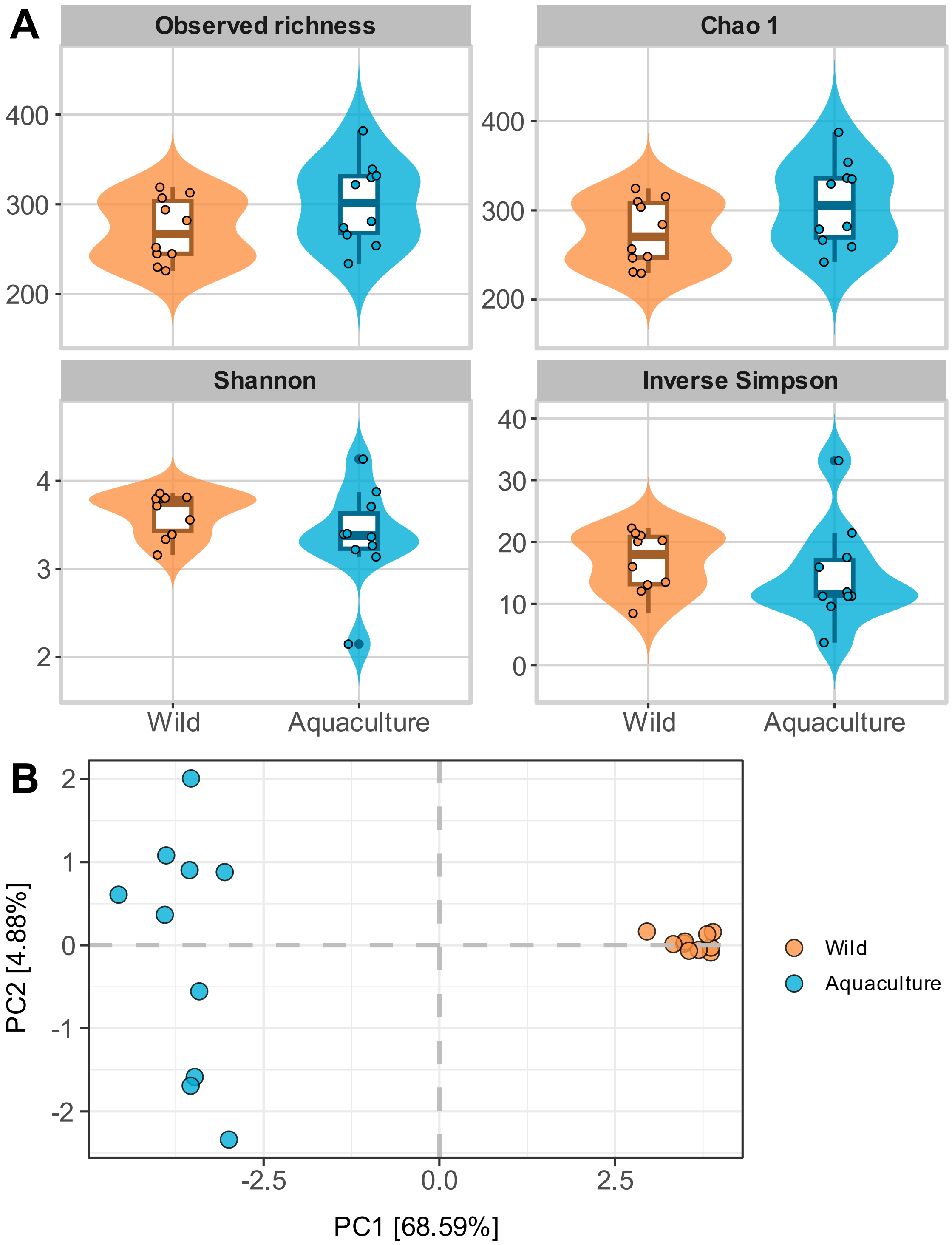


**Figure S4.** Diversity analysis at the genus level. (A) Alpha diversity measures for wild and aquaculture groups. Violin plots represents the alpha diversity indexes analyzed (Observed richness, Chao 1, Shannon and Inverse Simpson) in wild (orange) and aquaculture (blue) samples. (B) Beta diversity for wild and aquaculture groups. PCA plot is based on Aitchison distances at the genus level. Axes x and y represent the percentage of the total variance explained by the first two components, respectively.

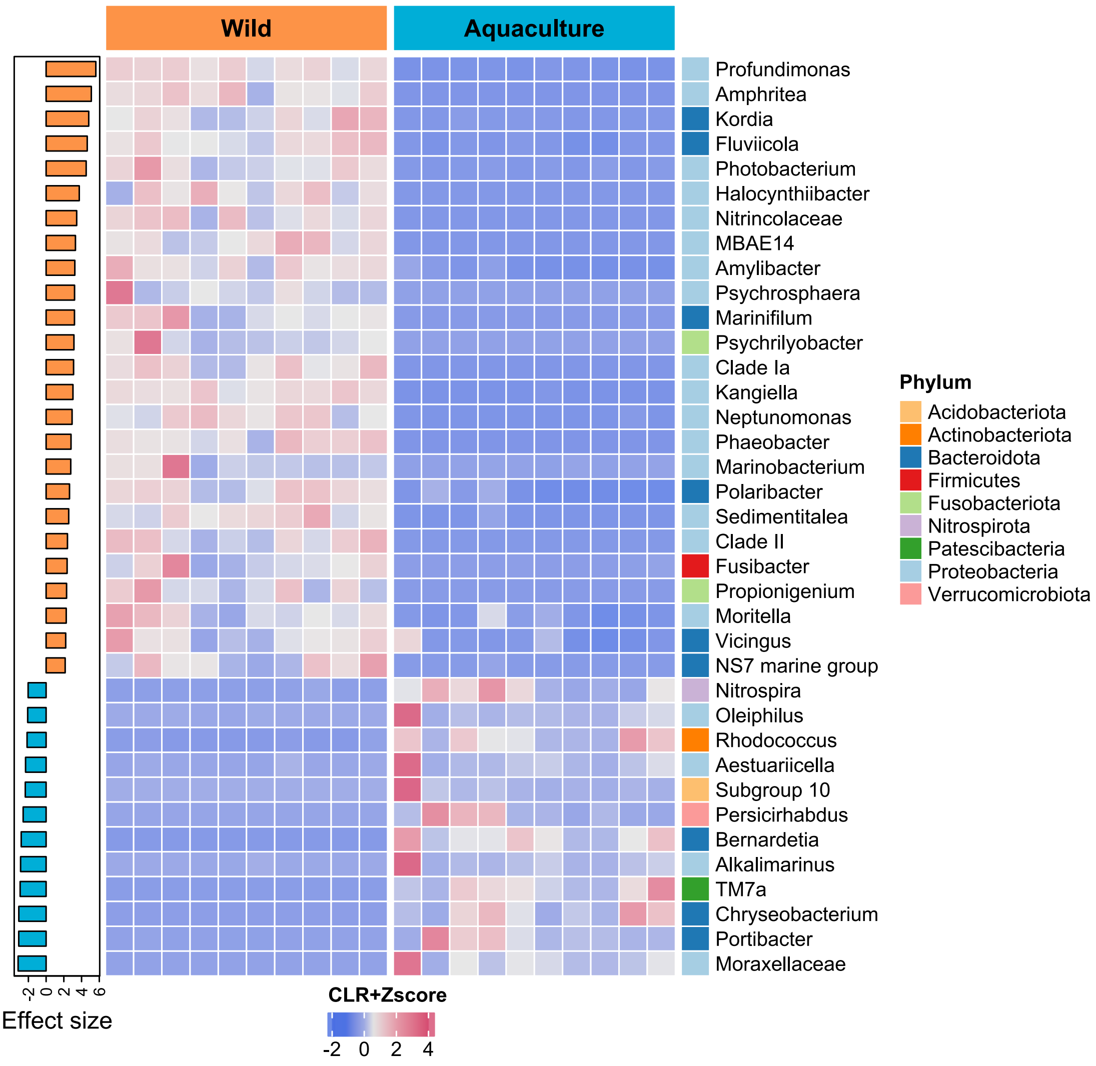


**Figure S5.** Differentially abundant genera between wild and aquaculture octopuses. The heatmap colors show the Z-scored CLR-transformed relative abundances of the 37 genera identified as significantly differentially abundant (FDR adjusted $p \leq 0.05$ and $|\text{effect size}| \geq 2.0$). The samples are ordered by hierarchical clustering in columns, while the bacterial species are sorted by decreasing effect size in rows, which is illustrated through a side barplot. The right sidebar displays the phyla to which each genus belongs.
